# Lysosomal acid lipase-activity as a novel target to efficiently address triple-negative breast cancer high malignancy

**DOI:** 10.1101/2023.10.01.560038

**Authors:** H. Steigerwald, T. Bozzetti, M. Tams, J. On, G. Hoffmann, J. Lambertz, K. Weidele, S. Treitschke, F. Reinhard, A. Kulik, N. Krawczyk, D. Niederacher, H. Neubauer, C. Werno, T. Rau, T. Fehm, K Esser

**Affiliations:** Heinrich-Heine-University of Düsseldorf, Department of Obstetrics and Gynecology, Düsseldorf, Germany; Center for Integrated Oncology (CIO Aachen, Bonn, Cologne, Düsseldorf), Germany; Medical Faculty, Institute of Anatomy and Cell-Biology, University of Bonn, Bonn, Germany; Fraunhofer Institute for Toxicology and Experimental Medicine ITEM, Regensburg, Germany; Institute of Pathology, University Hospital and Heinrich-Heine-University of Düsseldorf, Düsseldorf, Germany

## Abstract

Increased metabolism of neutral lipids, e.g. triglycerides and cholesterol esters, is a hallmark of malignant cancers such as triple-negative breast cancer (TNBC). Predominantly, cancer cells with a high epigenetic stem cell-associated signature increasingly utilize neutral lipids to maintain their high degree of tumor stemness, linking metabolic aberrations to epigenetically dysregulated differentiation processes. Lysosomal acid lipase (LIPA) is a central enzyme in the cellular utilization of exogenous and endogenous neutral lipids; however, the role of LIPA-activity in TNBC remains unexplored. We here show for the first time that pharmacological inhibition of LIPA, highly expressed in TNBC, reduces the expression markers of breast cancer stemness in cell culture models of TNBC. A role of LIPA in maintaining TNBC high cellular stemness was stressed by specific siRNA knock-down. Furthermore, inhibition of LIPA sensitized TNBC cells to therapy with Paclitaxel and Doxorubicin, two important chemotherapeutics in current TNBC treatment. When LIPA-activity was inhibited in a three-dinensional (3D) patient derived organoid model, we observed a significant reduction in TNBC cellular viability. Importantly, LIPA inhibition prevented tumor metastasis in a TNBC-zebrafish xenograft model *in vivo*. These findings introduce LIPA-activity as a novel pharmacological target in TNBC therapy to specifically address its high cancer malignancy with a potential for implementation of LIPA inhibitors into personalized treatment in the future.

## Introduction

Triple-negative breast cancer (TNBC) occurs in approximately 15% of breast cancer (BC). Treatment options are limited due to the missing expression of both hormone receptors (estrogen and progesterone receptor) and the HER2 receptor. Particularly, owing to its high aggressiveness, this BC subtype is characterized by an increased therapy resistance, disease recurrence, and early metastasis leading to the worst prognosis among BC intrinsic subtypes^1^. Additionally, a high degree of premenopausal women (<40 years) are diagnosed with TNBC. Consequently, there is an urgent medical need for new targeted therapies against this disease^2^.

It has become increasingly obvious that the epigenetically controlled low cellular differentiation status and high cancer cell stemness phenotypes are the main drivers for TNBC high malignancy^3^. In patients and cell culture models, BC stemness has commonly been characterized by the presence of cancer stem cells (CSC) with high expression patterns of tumor stem cell markers such as cluster of differentiation 44 (CD44^high^), aldehyde dehydrogenase activity (ALDH1^high^) and with primarily increased expressed in BRCA1 (breast cancer gene 1) mutated BC stem cells, SRY-Box transcription factor 2^high^ (SOX2 ^high^)^4^. Other proteins such as CD24 are expressed at low levels contributing to high CD44/CD24 ratio, which is characteristic for cells with stem cell features. These cells mainly contribute to drug resistance, metastasis, and disease relapse^5^. Therefore, targeting TNBC stemness has been regarded as a promising therapeutic approach to greatly improve current therapy and to realize personalized cancer treatment. Although the high cellular stemness is an important molecular characteristic of TNBC, there is limited information about the underlying pathophysiological dysregulations of the tumor cells to date.

The interface of cellular metabolism and epigenetic regulation mechanisms is a recently emerging area in cancer research since the enhanced metabolization of neutral lipids, such as triglycerides and cholesterol esters, has been demonstrated to be a key player in driving and maintaining cancer stemness^6–9^. Furthermore, a dysregulated utilization of neutral lipids is a hallmark of TNBC malignancy^10,11^. Notably, lysosomal acid lipase (LIPA) plays a central role in the cellular utilization of exogenous and endogenous neutral lipids^12^. However, the role of its enzymatic activity in TNBC malignancy has not been studied to date. Therefore, this study aimed to investigate the potential impact of LIPA’s enzymatic activity on the maintenance of TNBC cancer stemness and, linked to this pathophysiological process, on chemoresistance, proliferation, cellular viability and metastasis.

## Results

### LIPA is highly expressed in TNBC

Although the dysregulated neutral lipid metabolism in TNBC has been widely studied and LIPA enzymatic activity is essential for cellular utilization of neutral lipids, little is known about its function in TNBC. In accordance with a recently published study^13^ we detected high LIPA expression in TNBC tissue samples (immune reactive score: 10 +/-3.4) (Figure 1 A, B).

**Figure 1:**
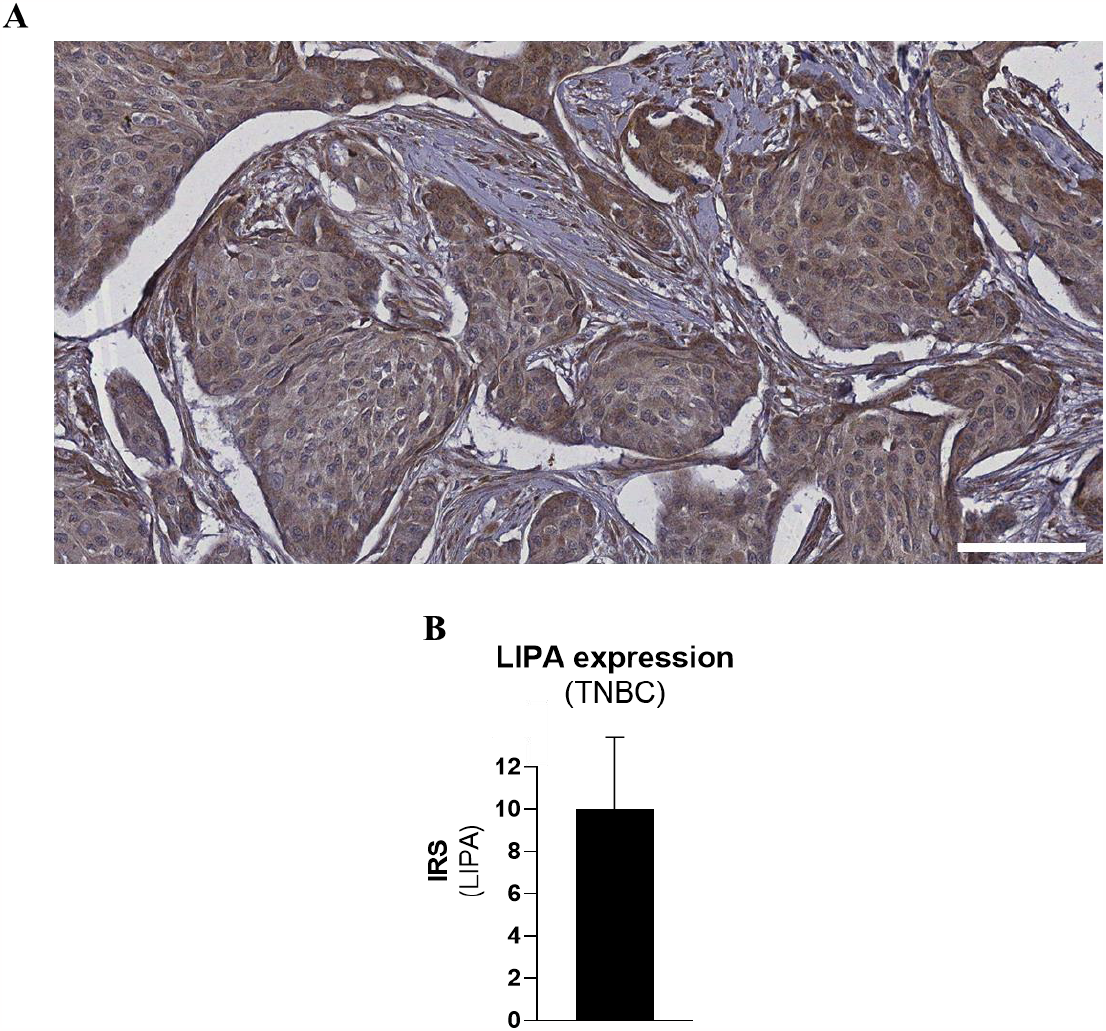
Immunohistochemical analysis of LIPA expression in TNBC. **(A, B)** TNBC tissues were paraffin fixed and tissue sections were stained with a LIPA-specific polyclonal antibody. (B) Immunreactive score (IRS) multiplicating intensity (0-3) with portion of positive cells (0: negative; 1: 10%; 2: 10-50%; 3: 51-80%; 4: >80%) yielding in a maximum of 12 was determined for TNBC tissue for 10 individual patients’ samples. *Scale is 50 μm.

### LIPA activity affects tumor neutral lipid metabolism and correlates with the expression of breast cancer stem cell markers

Since neutral lipid metabolism reportedly increases cancer malignancy including tumor stemness, we studied if LIPA-activity supports TNBC high stemness. The LIPA-specific inhibitor Lalistat was used to pharmacologically modulate LIPA enzyme activity in the TNBC cell lines MDA-MB-231 and MDA-MB-436. Cells were incubated with Lalistat for six days, and effects of LIPA-activity on cellular neutral lipid metabolism was analyzed by staining with BODIPY_493-503_. As illustrated in Figure 2a, LIPA inhibition strongly enhanced cellular neutral lipid content in MDA-MB-231 cells. Since LIPA inhibition leads to lysosomal and cytosolic accumulation of neutral lipids in a broad range of mammalian cells, this observation supports our hypothesis that LIPA activity plays an important role also in TNBC neutral lipid metabolism. Next, the effects on the expression of tumor stem cell-associated transcripts after LIPA inhibition were measured using reverse transcription-polymerase chain reaction (RT-PCR). In details, transcript levels of CD44, CD24 and additionally ALDH1 in MDA-MB-231 or SOX2 in BRCA1 mutated MDA-MB-436 cells were determined. As shown in Figure 2 B and C, mRNA expression levels of CD44, ALDH1 and Sox2 significantly decreased in TNBC-cells after Lalistat treatment. The expression ratio of CD44/CD24 also decreased significantly.

**Figure 2:**
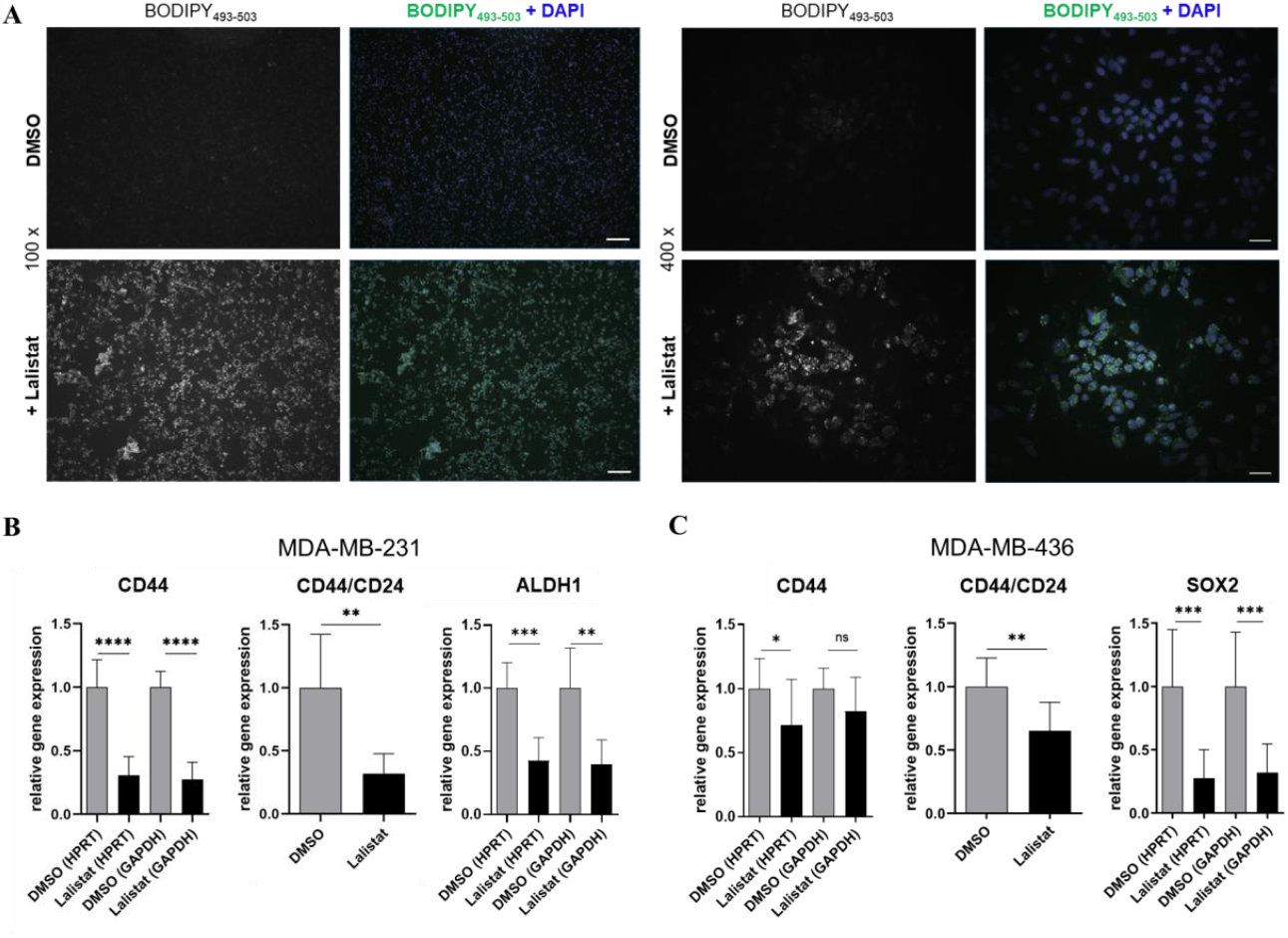
Lalistat affects tumor cell neutral lipid metabolism and reduces expression of tumor stem cell associated transcripts. *MDA-MB-231 (A, B) or MDA-MB-436 (C) cells were treated with 25 μM Lalistat or DMSO for six days. In (A) cells were fixed and stained for neutral lipids with BODIPY_493-503_. In (B, C) cells were lysed and the expression of the tumor stem cell markers CD44 (B, C), ALDH1 (B) and SOX 2 (C) was measured via qRT-PCR. The ratio of CD44/CD24 is also given (B, C). Indicated are mean values +/-SD, n=6 (B), n=9 (C). All obtained values were normalized to HPRT or GAPDH used as housekeeping genes. The values of relative gene expression were calculated using the 2 ^(-ΔΔCt)^– method. Statistical analysis was performed by using student’s t-test. * = p ≤ 0.05; ** = p ≤ 0.01; *** = p ≤ 0.001; **** = p ≤ 0.0001; ns: p = 0.11

Next, we investigated if the changes in stemness-associated transcripts detected by RT-PCR after LIPA-inhibition were correlated with analogous changes in protein expression. Flow cytometry analysis was used to investigate expression of stem cell markers CD44 and CD24 on protein level after treating MDA-MB-231 and MDA-MB-436 cells with Lalistat. As shown in Figure 3A and B, a significant reduction in the CD44 expression was detected in both cell lines, confirming that in TNBC cells the observed decreased CD44 mRNA can be translated into a reduced CD44 protein expression. Additionally, like observed on mRNA level, CD44/CD24-ratio was reduced in the MDA-MB-231-cells. To further validate the role of LIPA in the maintenance of TNBC stemness we performed siRNA-mediated specific LIPA knock-down leading to a residual LIPA-activity in cell lysates of about 7% and a significant reduction of the stem cell associated CD44 protein expression. Comparable like observed for Lalistat, CD44/CD24 protein expression ratio was also reduced (Figure S1).

**Figure 3:**
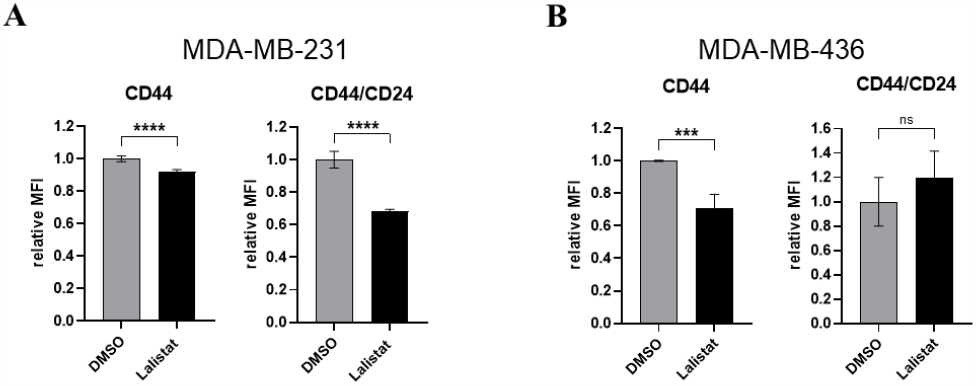
Pharmacologic inhibition of LIPA reduces CD44 protein expression. **(A, B)** Flow cytometry analysis of the change of CD44 protein expression and CD44/CD24 protein expression ratio in (A) MDA-MB-231 and (B) MDA-MB-436 cells after treatment with Lalistat. Cells were incubated with 25 μM Lalistat and corresponding DMSO control for six days and mean fluorescence intensity was measured after staining cells with CD44 and CD24 antibodies. The CD44 expression and the ratio of CD44/CD24 was set in relation to DMSO. Boxplots show the mean + SD, n=4. Statistical analysis was performed by using student’s t-test. *** = p ≤ 0.001; **** = p ≤ 0.0001; ns: p=0.24

### LIPA and tumor stem cell markers are co-expressed in breast cancer tissue

Since TNBC among BC is characterized by an extraordinary high cellular cancer stemness and according to our findings, that reduction of LIPA activity correlates with reduced expression of markers for cancer stemness in TNBC, we further analyzed the correlation of LIPA and tumor stemness marker expression levels in primary BC tissues using the database METABRIC in “cBioPortal for Cancer Genomics” ^14–17^. LIPA expression was confirmed to be associated with the mRNA expression of tumor stem cell markers ALDH1, CD44 and a high CD44/CD24 mRNA expression ratio (Figure 4A). In contrast, the expression levels of other cellular lipases which are reported to drive BC malignancy, such as hormone-sensitive lipase (HSL)^18^ and monoacylglycerol lipase (MGLL)^19^ or lipases exhibiting a tumor suppressor function like adipose triglyceride lipase (ATGL)^20^ are either not reduced or are much lower correlated with the expression of tumor stem cell markers (Figure 4B-D).

**Figure 4:**
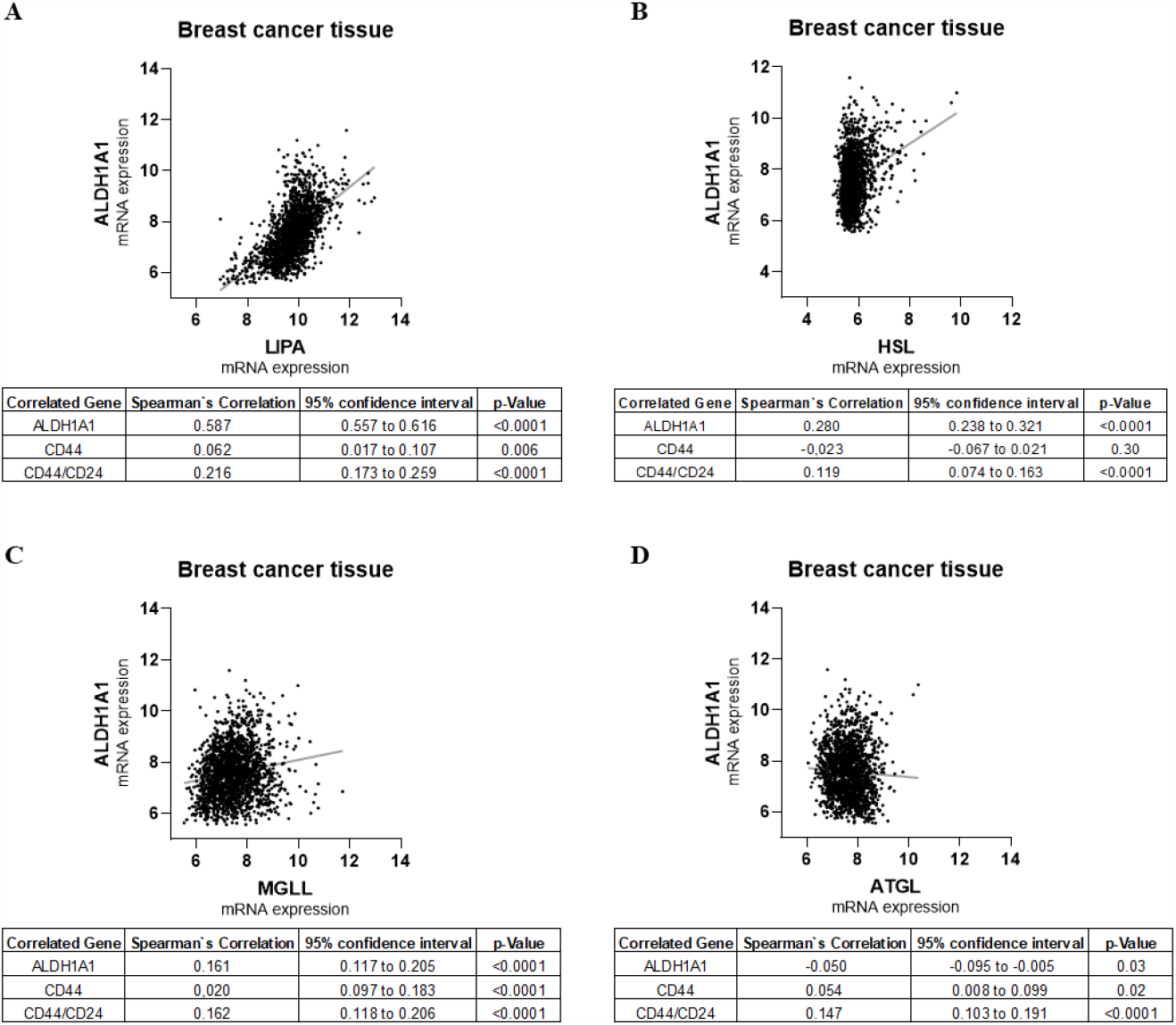
LIPA and tumor stem cell markers are co-expressed in breast cancer tissue. The co-expression of the cellular lipases LIPA (A), HSL (B), MGLL (C) and ATGL (D) with the tumor stem cell marker ALDH1A1, CD44 and the CD44/CD24 ratio were analyzed in tissue of breast cancer patients using the METABRIC database in “cBioPortal for Cancer Genomics”^14–17^. Scatter plots of the co-expression of lipases and ALDH1A1 as well as tables of co-expression data of lipases and ALDH1A, CD44 and CD44/CD24 expression, respectively, are shown.

### Lalistat inhibits cellular growth of MDA-MB-231 cells

In addition to driving and maintaining tumor cell stemness, the intratumoral utilization of neutral lipids is closely associated with proliferation of breast cancer cells. We therefore investigated if TNBC cell proliferation was also affected by treatment with LIPA inhibitor Lalistat.

As shown in Figure 5A, a significant reduction in cell proliferation up to 32% was observed in MDA-MB-231 cells using Lalistat concentrations from 12.5 – 50 μM. To exclude toxic effects also leading to reduced cellular ATP levels determined in the cell proliferation assay, cellular toxicity of Lalistat at different concentrations was excluded by CellTox™ Green Cytotoxicity Assay (Figure S2). In contrast to MDA-MB-231 cells a slight but significant anti-proliferative effect was observed in MDA-MB-436 cells only at 50 μM Lalistat (Figure 5B).

**Figure 5:**
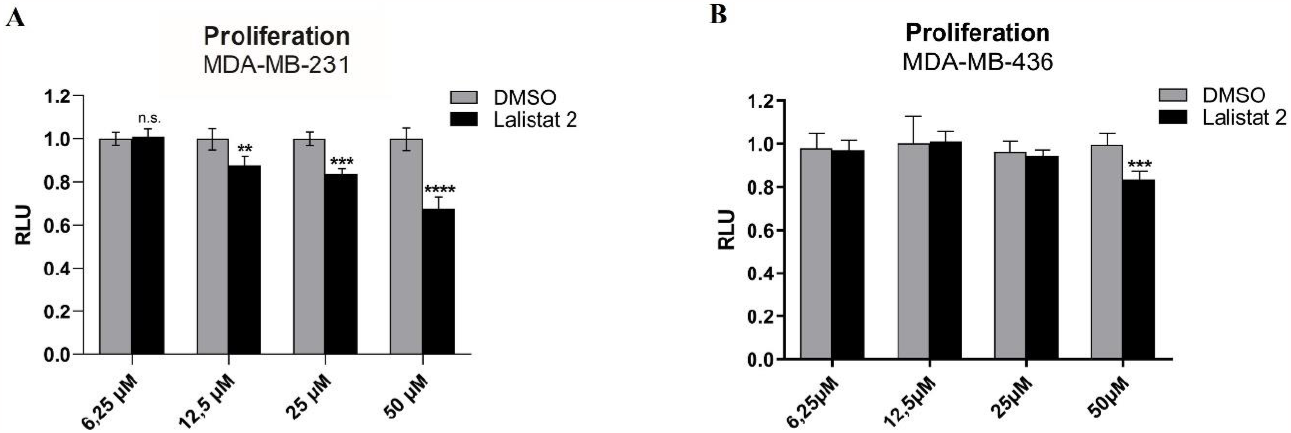
Lalistat inhibits cellular growth of MDA-MB-231 cells. MDA-MB-231-cells **(A)** and MDA-MB-436-cells **(B)** were treated with Lalistat in dilution series from 50 μM to 6,25 μM for six days and cellular ATP was measured by CellTiter-Glo (RLU = relative luminescence). The graph shows the mean + SD, n=4 (A) and n=12 (B). All values were set in relation to DMSO. Statistical analysis was performed by using one-way Anova. ** = p ≤ 0.01; *** = p ≤ 0.001; **** = p ≤ 0.0001

### Lalistat sensitizes TNBC to chemotherapy

Deregulated cellular neutral lipid metabolism has been reported to contribute to BC stemness and reduce chemosensitivity^6,8^. To explore the hypothesis that Lalistat sensitizes TNBC to cell death-inducing effects of chemotherapeutics, MDA-MB-231 and MDA-MB-436 cells were incubated with Lalistat in combination with paclitaxel in dilution series. As shown in Figure 6A, Lalistat dose dependently induced an over threefold reduction of the half-maximal inhibitory concentration (IC_50_) of paclitaxel in MDA-MB-231 cells. Similarly, Lalistat diminished the IC_50_ of paclitaxel in MDA-MB-436 cells up to 2.6-fold (Figure 6B). Interestingly, Lalistat significantly enhanced the effect on paclitaxel induced toxicity compared to Chloroquine, an agent already established in clinical studies for targeting TNBC stemness and increasing cancer chemosensitivity^21^ (Figure S3A).

**Figure 6:**
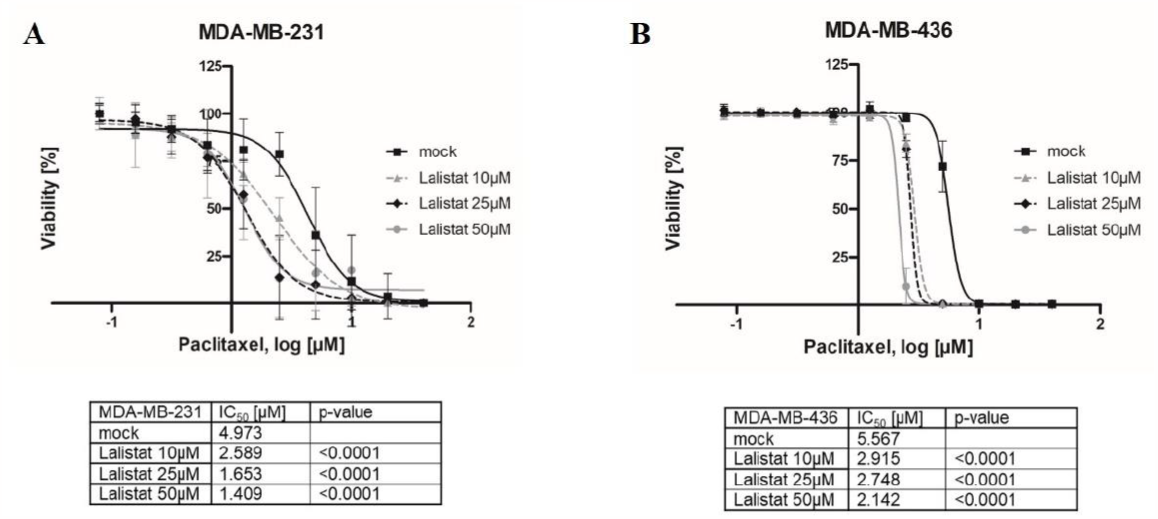
Lalistat sensitizes TNBC to anti-cancer activity of chemotherapeutics. **(A, B)** IC_50_-curves of paclitaxel in the absence or presence of different concentrations of Lalistat. MDA-MB-231 cells (A) and MDA-MB-436 cells (B) were pretreated with Lalistat at concentrations indicated for three days, followed by incubation for another three days with same Lalistat concentrations, respectively, and paclitaxel in dilution series. The viability of the cells was evaluated by CellTiter-Glo^®^. Depicted is the mean +/-SD, n=20 (A) and n=4 (B). Mock corresponds to DMSO 0.075% with no difference to medium alone. Tables listed below each graph summarize the calculated IC_50_ values. P-values were calculated relatively to mock treatment. All values were set in relation to viability of untreated cells. Statistical analysis was performed by using one-way Anova.

In addition to enhanced utilization of exogenous lipids, LIPA plays an important role in the hydrolysis of endogenous neutral lipids in a process called “lipophagy”, a specialized form of autophagy, which provides the cells with an alternative source of energy, especially under nutrient depletion^12,22^. Therefore, the effect of Lalistat on paclitaxel-induced cell death was investigated in a neutral lipid-free medium. As shown in Figure S3B, a significant reduction of about two-fold in the IC_50_ of paclitaxel compared to control MDA-MB-231 cells was detected. Thus, although the observed reduction in IC_50_ was significantly less, LIPA inhibition also led to a significant increase in paclitaxel-induced cell death in the absence of exogenous neutral lipids.

Since current chemotherapy of TNBC in addition to taxans includes the application of anthracyclines, we investigated if Lalistat also increases the anti-tumor effect of the anthracycline Doxorubicin. When MDA-MB-231 cells were incubated with Doxorubicin in the presence of Lalistat, a remarkable decrease of 53.3% in IC_50_ was observed (Figure S3C). This synergistic effect of Lalistat was also observed on proliferation at non-toxic concentrations of Doxorubicin (Figure S3D, E).

To ensure a potential therapeutic window and to rule out substantial toxicity in non-malignant cells, the IC_50_ of Lalistat was determined after treating the non-tumorigenic epithelial breast cell line, MCF 10A, resulting in an IC_50_ of 96.28 μM (Figure S3F). Thus, the concentrations used in these studies for TNBC cell line treatment were significantly lower than that necessary to induce cellular toxicity in non-malignant breast cells.

### Inhibition of LIPA-activity reduces cellular viability in a patient-derived TNBC organoid model

To reflect the 3D situation in a human tumor *in vivo* and to investigate if inhibition of LIPA also has an effect on primary TNBC cells, we used an organoid gained with tumor which had been established from tumor cells isolated from a biopsy of TNBC patient prior to neoadjuvant therapy (Figure 7A, B). Interestingly, in contrast to the 2D cell culture model, we observed a direct effect of LIPA inhibitor Lalistat on the cellular viability without the addition of a chemotherapeutic drug, which was confirmed by microscopic analysis showing a dose dependent loss of vital organoids (Figure 7A) and by measuring a significant reduction in cellular ATP levels (Figure 7B).

**Figure 7:**
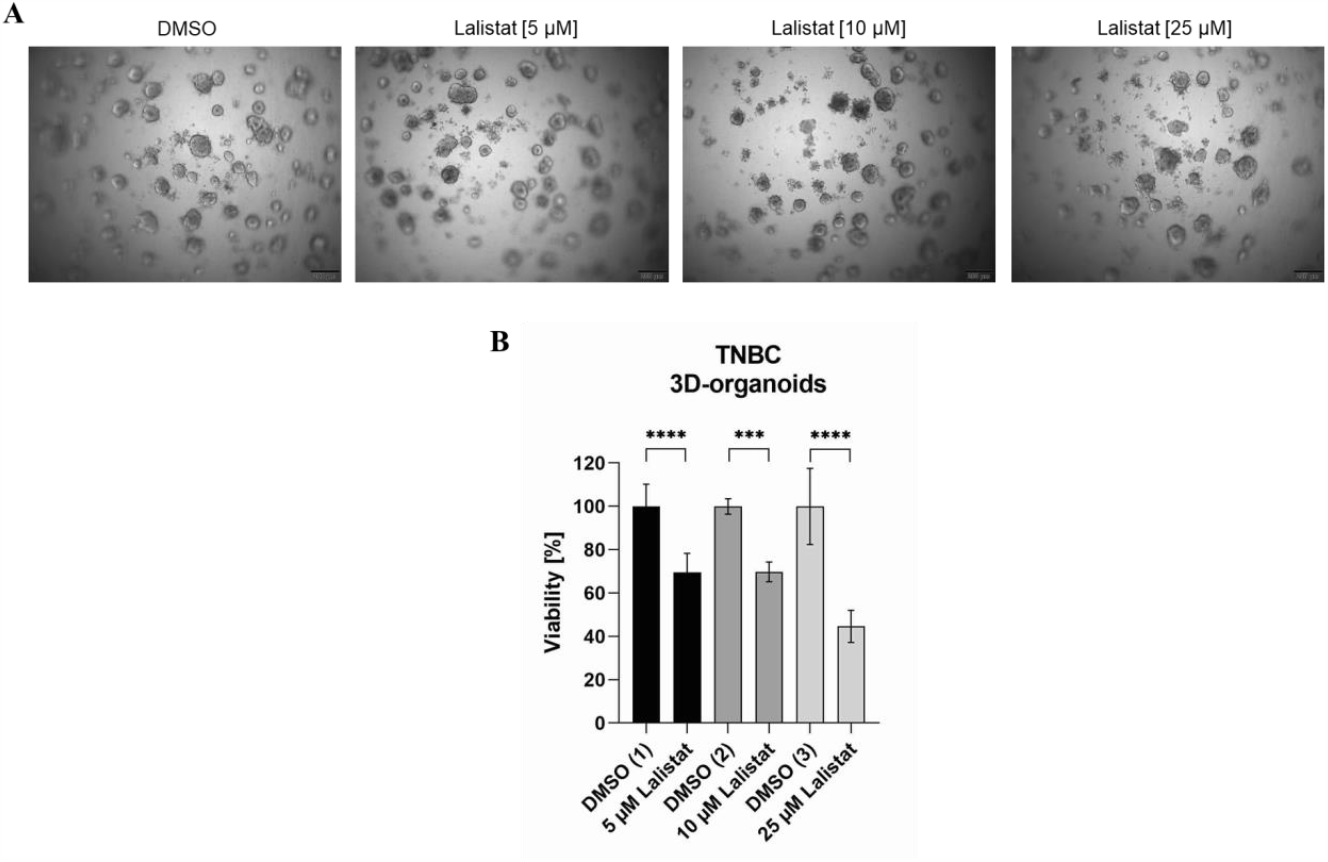
Lalistat reduces cellular viability in a patient-derived TNBC organoid model. **(A, B)** Patient-derived TNBC-cells were cultured as Organoids and incubated for 6 days with 5 μM, 10 μM and 25 μM Lalistat or DMSO as control. Subsequently, organoids were analyzed by brightfield fluorescence microsopy (A) and or cellular viability was investigated by CellTiter Glo® 3D (B). Scale bars are 200 μm. All values were set in relation to untreated cells. Statistical analysis was performed by using one-way Anova. *** = p ≤ 0.001; **** = p ≤ 0.0001.

### LIPA-inhibition diminishes MDA-MB-231 metastasis in vivo

Since early metastasis is a hallmark of TNBC and high cancer stemness is regarded to represent a main driver of this process, we studied the effect of LIPA inhibition on MDA-MB-231-tdTomato cell metastasis in a Zebrafish xenograft model *in vivo*. As shown in Figure 8A, pretreatment of injected tumor cells with Lalistat inhibited spread of MDA-MB-231-tdTomato cells into the head and tail, respectively, leading to a 4.7-fold reduction of metastasis over DMSO-treated tumor cells (Figure 8B).

**Figure 8:**
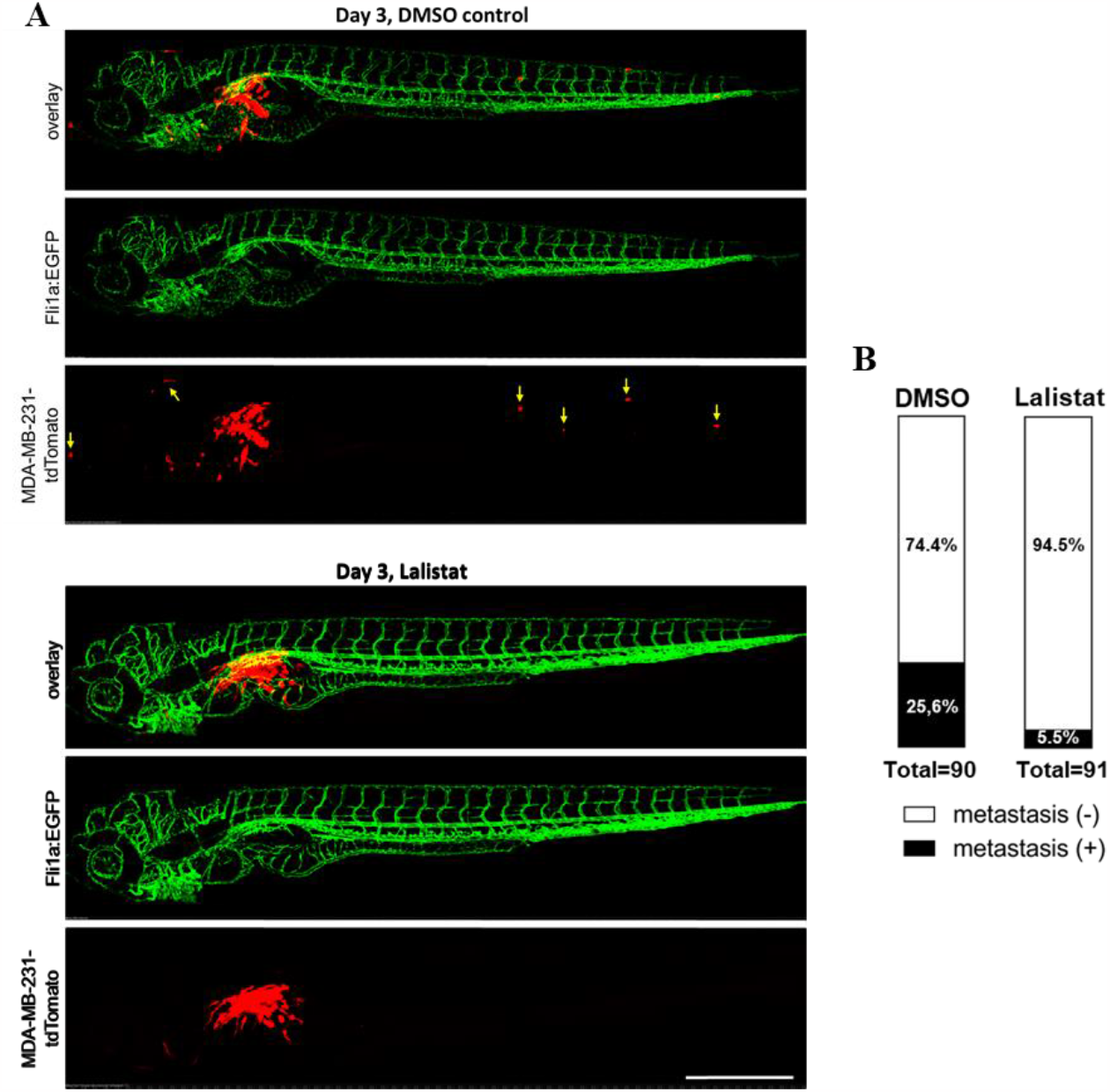
Lalistat inhibits tumor metastasis in an MDA-MB-231-tdTomato zebrafish xenograft model. **(A, B)** MDA-MB-231-tdTomato were preincubated for 6 days with 25 μM Lalistat or DMSO as control. 200 MDA-MB-231-tdTomato were injected into perivitelline space of 3 days old zebrafish larvae (181 fishes in total of three independent experiments). After two days zebrafishes were analyzed by confocal microscopy (A; one representative zebrafish each) or brightfield fluorescence microscopy (B) and fishes positive or negative for tumor metastasis were counted (B). Arrows in (A, upper picture) depict cancer cells metastasized into zebrafish head and tail, respectively. Statistical analysis of difference in zebrafishes positive for metastasis between DMSO and Lalistat treated tumor cells (B) was performed using Chi-square test: p_(DMSO/Lalistat)_=0.00016

## Discussion

To develop an effective and personalized therapy against aggressive cancers like TNBC, it is important to detect novel cellular targets^23^. One important candidate field is epigenetic aberrations which, in addition to genetic mutations, should function as a major driver for cancer malignancy^24^. TNBC cells are characterized by a low differentiation status, indicating wide epigenetic dysregulations in physiological differentiation processes. These cancer dysregulated mechanisms enhance cellular tumorigenicity, increase resistance to apoptosis and reduce cellular immunogenicity. Thus, moving cancer to a higher differentiation status and lower cellular stemness stage may be a promising strategy to improve current TNBC therapy^3,25,26^. However, the available epigenetically acting drugs like histone deacetylases, DNA-methyltransferases, or bromodomain and extra-terminal protein inhibitors address ubiquitous epigenetic regulators, controlling a wide range of physiological processes, including cancer promotion and suppression, characterized by only limited cancer pathology specificity^27,28^.

This study is the first to reveal that LIPA-activity correlates with the expression of breast cancer stem cell markers in cell culture models of TNBC supporting a central role in maintaining high tumor stemness in this breast cancer subtype. In line with this hypothesis, LIPA inhibition reduces expression of cancer stem cell markers but increases sensitization to chemotherapy and reduces metastasis *in vivo*. Since our bioinformatics analysis of primary human BC tissue emphasizes a strong correlation of LIPA expression with high cancer stemness markers in BC patients and LIPA is highly expressed in primary TNBC tissue in our study, pharmacological inhibition of LIPA may efficiently be implemented into future personalized (combined) therapy regimens of TNBC.

Although metabolic aberrations in highly malignant and tumorigenic cancers like TNBC have been widely reported, their impact on cellular stemness and the key factors involved are only partially understood. Notably, cancer cells utilize free fatty acids for mitochondrial β-oxidation which have been shown to significantly drive cancer stemness. Furthermore, neutral lipids fill up cellular storages (“lipid droplets”), constitute precursors for signaling molecules and build up elements of cellular membranes, in particular lipid rafts^6–8^. These processes are widely described to drive and maintain cancer cell stemness and likely represent main mechanisms how LIPA contribute to TNBC high malignancy.

Interestingly, in contrast to the studies performed in 2D cell culture models we observed a direct effect of pharmacological LIPA inhibition on cellular viability in a patient-derived TNBC organoid model underlining its high therapeutic potential in humans. Actually, it occurs reasonable that in a more physiological 3D tumor architecture interference with the TNBC-characteristically dysregulated neutral lipid metabolism illustrates a superior anti-cancer activity compared to the 2D situation since it has been well described that neutral lipid metabolism including fatty acid oxidation is highly increased in 3D compared to 2D cancer models ^29^. Thus, high metabolization of neutral lipids might more contribute to tumor cells’ abilities to compensate apoptosis inducing cellular stress signals compared to 2D cultured cells. This is probably even more relevant in the rather hypoxic inside of the organoids^29,30^.

To date, only limited investigations have been performed to target enhanced neutral lipid utilization in cancer. Orlistat, a broad inhibitor of neutral lipid hydrolysis, exhibits anticancer activity. However, this anti-tumor function has been attributed to inhibiting fatty acid synthase but not to targeting neutral lipid hydrolyzing lipases^31^. Distinct cellular lipase activity, as described for ATGL, has been shown to mediate anticancer function^20,32,33^. Thus, besides leading to severe side effects, targeting a broad range of cellular neutral lipid lipases appears unsuitable for effective cancer therapy. Similarly, strategies to target enhanced uptake of exogenous neutral lipids in cancer by addressing different kinds of receptors of the low-density lipoprotein (LDL)-receptor family class, such as LDL receptor, LDL receptor-related protein, megalin, and very low-LDL receptor, might not be efficient since these receptors can partially compensate each other if one or more are not fully functional^34–36^. Furthermore, since cancer pathology includes the utilization of exogenous and endogenous lipids, this concept does not fulfill the dual potential role in anticancer therapy as described for LIPA-inhibition.

Importantly, when this manuscript was composed, Liu et al. was published showing LIPA as a potential target in TNBC therapy^13^. The authors found a so far undescribed chaperone function of LIPA in the endoplasmatic reticulum, which was independent of its enzymatic activity. In concordance with our studies the authors did not observe cell toxicity when inhibiting LIPA activity in 2D TNBC cell lines. In contrast to our study Liu and colleagues did not investigate a potential effect of LIPA activity on cellular high tumor stemness in TNBC. Based on results from our studies we conclude that, in addition to its newly defined chaperone function, LIPA activity plays an important role in TNBC pathophysiology and significantly contributes to TNBC high stemness phenotype by generating free fatty acids and free cholesterol via its neutral lipid hydrolysing activity. This newly described function should be taken into account when investigating LIPA as a potential drug target in future TNBC treatment. Since TNBC is divided into different molecular subtypes this might especially be important in the mesenchymal stem-like subtype illustrated to possess an extraordinarily high tumor stemness among TNBC^37^. This was also considered in the studies presented here with using MDA-MB-231 and MDA-MB-436 cells as cellular models of the mesenchymal stem-like TNBC subtype^37^.

Taken together, this study is the first to describe that pharmacological inhibition of LIPA, highly expressed in TNBC, reduces cancer stemness and sensitizes TNBC cells to cell death e.g. induced by chemotherapy as well as diminishes their metastasis. According to our current model (Figure 9), the underlying pharmacological mechanism likely includes the inhibition of cancer cell utilization of neutral lipids, derived from extracellular sources via lipoprotein endocytosis or delivered intracellularly from endogenous sources via lipophagy. The reduced cellular disposing ability for free fatty acids and cholesterol by LIPA inhibition resulted in decreased free fatty acid oxidation by the mitochondria, filled cellular lipid droplets, and the generation of lipid rafts and mediators, which led to diminished cellular stemness. Further *in vivo* investigations are underway to evaluate the therapeutic potency of pharmacological LIPA inhibition to specifically address TNBC tumor malignancy.

**Figure 9:**
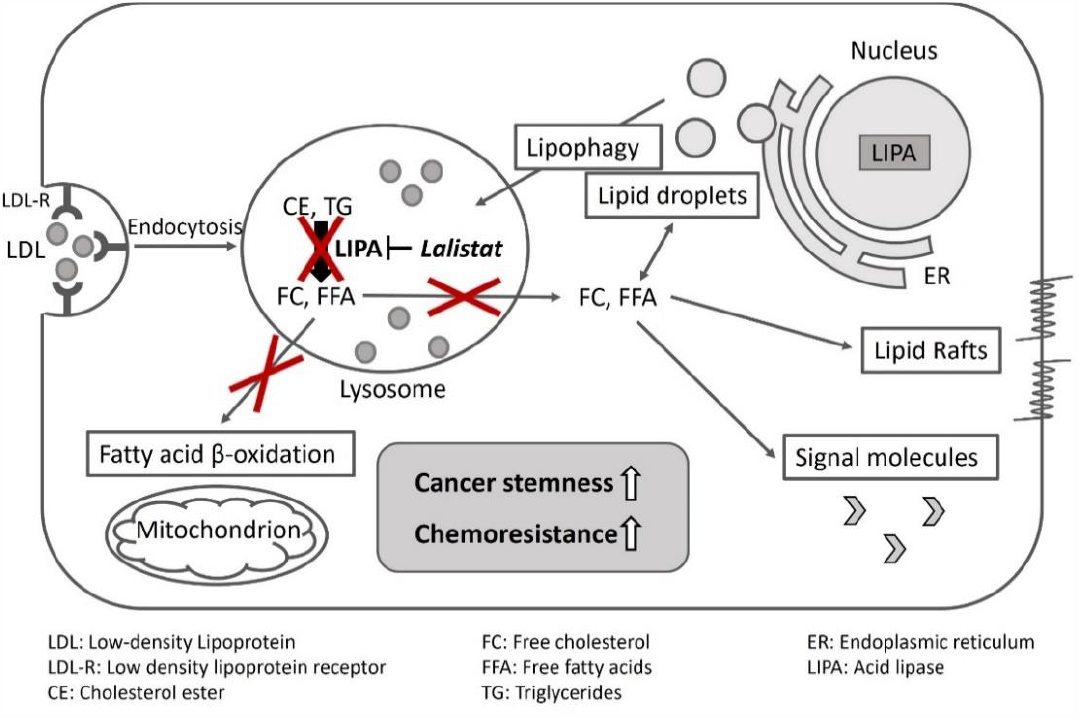
Model for LIPA inhibition and reduction of TNBC high malignancy.

## Materials and methods

### Cell culture

All cell lines were purchased from ATCC (LGC Standards GmbH, Wesel, Deutschland). MDA-MB-231-tdTomato were a gift of Arwin Groenewoud and Felix Engel, Univeristätsklinikum Erlangen, and generated by transduction of MDA-MB-231 with a pLenti-V6.3 Ultra lentiviral CMV driven tdTomato-P2A-T2A, Blasticidin selectable. MDA-MB-231 (ATCC®HTB-26™) and MDA-MB-436 (ATCC®HTB-130™) were cultured with Roswell park memorial institute (RPMI) Medium 1640 supplemented with 15% fetal bovine serum and 1% penicillin/streptomycin (Gibco, USA) and incubated at 37°C in 5% CO_2_. Non-tumorigenic MCF-10A cells (ATCC^®^ CRL-10317^™^) were cultured with Dulbeccos’s modified Eagle medium (DMEM) F12 (Gibco, UK) supplemented with 5% horse serum (Gibco, USA), 1% penicillin/streptomycin (Gibco, USA), epidermal growth factor (EGF) at 20 ng/ml (Peprotech), hydrocortisone 0.5 mg/ml (Sigma, #H-0888), cholera toxin 100 ng/ml (Sigma, #C-8052) and insulin 10 μg/ml (Sigma, #I-1882). Incubation took place at 37°C in 5% CO_2_.

### Expression analyses of tumor stem cell markers

Cells were incubated for six days with Lalistat (Lalistat 2, TOCRIS, Bristol, UK) in a 6-well plate. mRNA was isolated using ReliaPrepTM RNA Cell Miniprep System (Promega, USA), and cDNA synthesis was performed using Superscript® First-Strand Synthesis System for RT-PCR (Thermo Fischer Scientific, Carlsbad, USA) according to the manufacturer’s protocols. For gene expression analysis, appropriate primers (metabion international AG, Germany) and LightCycler^®^ 480 SYBR^®^ Green I Master (Roche, Mannheim, Germany) were used. RT-PCR was performed using LightCycler® 480 Instrument II (Roche, Germany).

### Knock-down of acid lipase

Seeded cells were treated with Lipofectamine^®^ RNAiMAX Reagent (Invitrogen by Thermo Fischer Scientific, Carlsbad, USA) according to the manufacturer’s protocol. A target-specific Silencer^®^ Select siRNA (Thermo Fisher Order Number: 4392420; siRNA ID: s8199) and a corresponding control siRNA (Thermo Fisher Order number: 4390843) were used. Knock-down was performed in serum-free Advanced RPMI 1640 Medium (Gibco, UK) before cells were further cultured, as mentioned in section “cell culture”.

### Immunhistochemical analysis of TNBC tissue

Whole tissue sections were obtained from routine formalin-fixed paraffin-embedded (FFPE) blocks of TNBC patients who underwent surgery or were tissue biopsied between the years 2007 and 2022 at the Department of Obstetrics and Gynecology, Heinrich-Heine-University of Düsseldorf. All tumor samples were obtained with informed patients’ consent and the pathology files retrieved as approved by the Ethics Committee of the Medical Department of the Heinrich-Heine-University of Düsseldorf (study number: 2021-1589).

Tissue sections were deparaffinized and rehydrated by washing with Xylene following alcohol grade steps from 100% to 50% Ethanol and deionized water. Antigen retrieval was performed in 10 mM sodium citrate buffer (pH 6.0). Endogenous peroxidas activity was quenched with 3.0% hydrogen peroxide for 15 min, tissue was permeabolized with 0,5% Triton X-100 in PBS (PBS-T) and non-specific binding was blocked by incubating the tissue section with blocking reagent (DAKO). Antibody staining was performed in PBS-T with 1% blocking reagent (DAKO) using 1/200 anti-LIPA (MyBioSource, MBS7004224) following 1/2000 secondary antibody. For detection, slides were incubated with 3,3′-Diaminobenzidine following Hematoxylin after washing.

### Western blot

After cell lysis using radioimmunoprecipitation assay (RIPA) buffer (ThermoFisher Scientific), 25 μg protein extracts were separated by sodium dodecyl sulfate (SDS) gel electrophoresis and transferred to a polyvinylidene difluoride (PVDF) membrane (BioRad) overnight. For LIPA detection, primary antibody (Abcam, ab154356) diluted 1:1000 in 0.5% milk powder Tris-buffered saline, 0.1% Tween 20 (TBST) and secondary anti-rabbit IgG (Cell Signaling, #7074S) were used. For STAT3 and pSTAT3, rabbit monoclonal antibodies 79D7 and D3A7 (Cell signaling) were diluted 1:1000 in 5% bovine serum albumin-TBST (BSA-TBST). β-actin and glyceraldehyde 3-phosphate dehydrogenase (GAPDH) expression were determined using mouse antibodies sc-47724 and sc-47778 (Santa Cruz) (1:2500) and anti-mouse IgG (Cell Signaling, #7076S), the secondary antibody. Analysis was performed by Image Lab software.

### LIPA activity assay

Cellular lipase activity assay was performed as described^38^. Briefly, 4-methylumbelliferone palmitate (4-MUP, Cayman Chemical) 4-MUP was dissolved in DMSO at 13.3 mM and added to 150 mM sodium acetate buffer pH 4.0 containing 1.0% (v/v) Triton X-100 and 0.5% (w/v) cardiolipin to get a 0.345 mM substrate solution. Cells were lysed in 150 mM sodium acetate buffer pH 4.0 with 1.0% (v/v) Triton X-100. For enzyme reaction, 40 μl of cell lysate and 10 μl DMSO or 30 μM Lalistat were combined and preincubated for 10 minutes at 37°C before 150 μl substrate solution was added. Samples were incubated at 37°C for 3h and subsequently measured in black 96-well plates in a fluorescence microplate reader (TECAN) at 320 nm (excitation) and 460 nm (emission). Reaction was stopped by adding 100 μl 15 mM HgCl2 to all sample wells. The LIPA enzyme activity was calculated by subtracting the enzymatic activity of the reaction inhibited by Lalistat from the inhibited reaction.

### Flow cytometry analysis

After treatment, cells were washed with Phosphate-buffered saline (PBS) and dissociated using Cell Dissociation Buffer (Gibco, 13151-014). After performing Fc block by using blocking buffer (Dako, X0909), cells were stained by CD24 (Abcam, SN3) and CD44 (Novus, MEM-263) antibodies diluted in antibody-diluent (Dako, S2022) according to the manufacturer’s protocol. Then, the cells were washed and taken up in the fluorescence-activated cell sorting (FACS) buffer consisting of PBS, 2 mM Ethylenediamine tetraacetic acid (EDTA) (Sigma, E6758) and 0.2% BSA (Sigma, A9418). After adding Propidium iodide (Gibco, P1304MP) for staining dead cells, samples were analyzed by flow cytometer. All steps were performed at 4°C.

### Bioinformatical analysis of breast cancer tissues

The data for co-expression of LIPA and the tumor stem cell markers ALDH1A1, CD44 and CD24, such as proteins associated with STAT3 signaling in the tissue of BC patients, were provided by the database “cBioPortal for Cancer Genomics”.

### Cell viability and toxicity assays

For effect on cell viability by Lalistat treatment alone, cells were treated for six days in the concentrations indicated. For investigation of synergistic effects by Lalistat, cells were pretreated with Lalistat for three days and incubated either for three days with a combination of Lalistat or Chloroquine (Sigma, Steinheim, Germany) and NeoTaxan® (Hexal, Holzkirchen, Germany) or for six days with a combination of Lalistat and Doxorubicin (Cayman Chemical, USA) in the concentrations indicated. Cell viability was determined using CellTiter-Glo® Luminescent Cell Viability Assay (Promega, Madison, USA) according to the manufacturer’s protocols. After incubation with the drugs mentioned, cells were lysed with the reaction lysis buffer and incubated for 10 minutes at room temperature. Subsequently, luminescence was measured in a microtiter plate reader.

To distinguish the toxic and anti-proliferative effects of Lalistat in DMA-MB-231 cells, the CellTox™ Green Cytotoxicity (Promega, Madison, USA) was used to determine cell toxicity according to the manufacturer’s protocol.

All measurements were performed using the Spark® multimode microplate reader (Tecan, Switzerland).

### Patient-derived TNBC organoid model

The patient-derived TNBC organoid model (cultured from a fine needle tumor biopsy sample taken before neoadjuvant chemotherapy) has been established and characterized at Fraunhofer ITEM Regensburg using published culture conditions^39^. Malignant origin was verified by detection of copy number alterations. TNBC origin was confirmed by a human pathologist after examination of Formalin-fixed, paraffin-embedded tissue sections immune-stained for ER, PR, HER2, and Ki67 according to routine laboratory staining protocol and using conventional light microscopy.

For drug testing (described in detail by Calandrini et al.^40^), organoids the size of 50-100 μm were harvested and resuspended in organoid culture medium supplemented with 5% Geltrex (Gibco). Using Multidrop Combi (Thermo Scientific) ∼400 organoids per well were plated on μclear 384-well plates (Greiner) precoated with 10 μl Geltrex. After 1h hour incubation, Lalistat was added using D300e Digital Dispenser (Tecan). Six days after treatment, morphology was visualized with Olympus IX81 microscope. Subsequently, CellTiter-Glo® 3D Cell Viability Assay (Promega) was performed according to the manufacturer’s instructions and luminescent signals were acquired with an EnVision plate reader (Perkin Elmer).

### Evaluation and statistical analysis

All analyses were performed using GraphPad PRISM 8.4.3, Graphpad Software, Inc. For statistical analysis, student’s t-test and ANOVA with Tukey’s multiple comparisons tests for group comparisons were used. A probability of error of p < 0.05 was considered statistically significant.

## Supplemental Data

**Figure S1:**
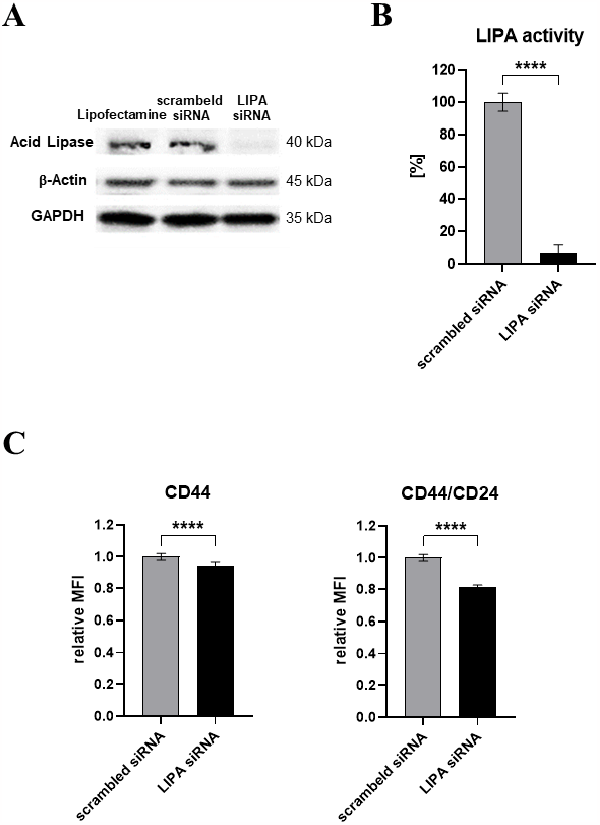
Gene knockdown of LIPA reduces CD44 protein expression. **(A-C)** Single cell analysis of the change in expression of CD44 in MDA-MB-231 after LIPA gene knockdown was performed. MDA-MB-231 cells were incubated with acid lipase-specific (LIPA) Silencer® Select siRNA or scrambled siRNA as negative control for three days before cells were cultured for further three days without siRNAs. In (A), Western Blot was performed using anti-LIPA antibody and GAPDH and ß-Actin antibody as loading control (left panel) or LIPA-activity was determined in cell lysates (right panel). Mean fluorescence intensity was measured after staining cells with CD44 and CD24 antibodies by flow cytometry analysis. The CD44 protein expression and CD44/CD24 protein expression ratio was set in relation to scrambled siRNA. The graph shows the mean + SD, n=4. Statistical analysis was performed by using student’s t-test. **** = p ≤ 0.0001

**Figure S2:**
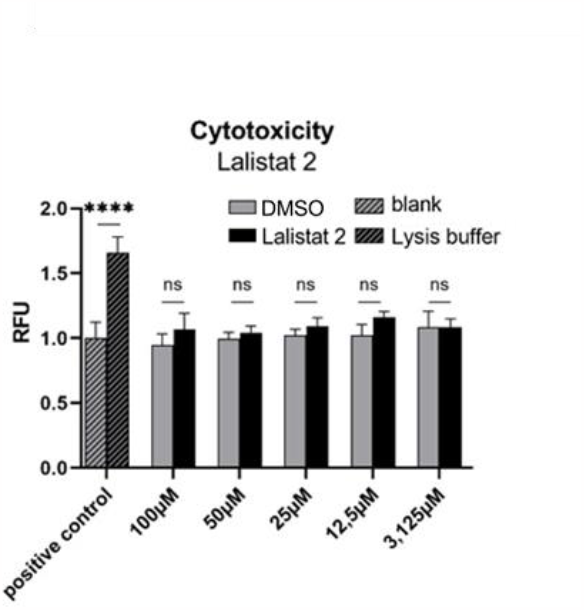
Lalistat is not toxic in MDA-MB-231. Toxicity of Lalistat in concentrations used was excluded by toxicity tests. MDA-MB-231 cells were treated with Lalistat in concentrations from 3,125 μM to 100 μM. The cell toxicity was determined in form of relative fluorescence by using CellTox™ Green Cytotoxicity Assay measuring relative fluorescence units set free from dead/lysed cells. The positive control corresponds to lysis solution included in the assay. The graph shows the mean + SD, n=4. All values received were set in relation to blank-mean in form of untreated cells. Statistical analysis was performed by using one-way Anova. **** = p ≤ 0.0001

**Figure S3:**
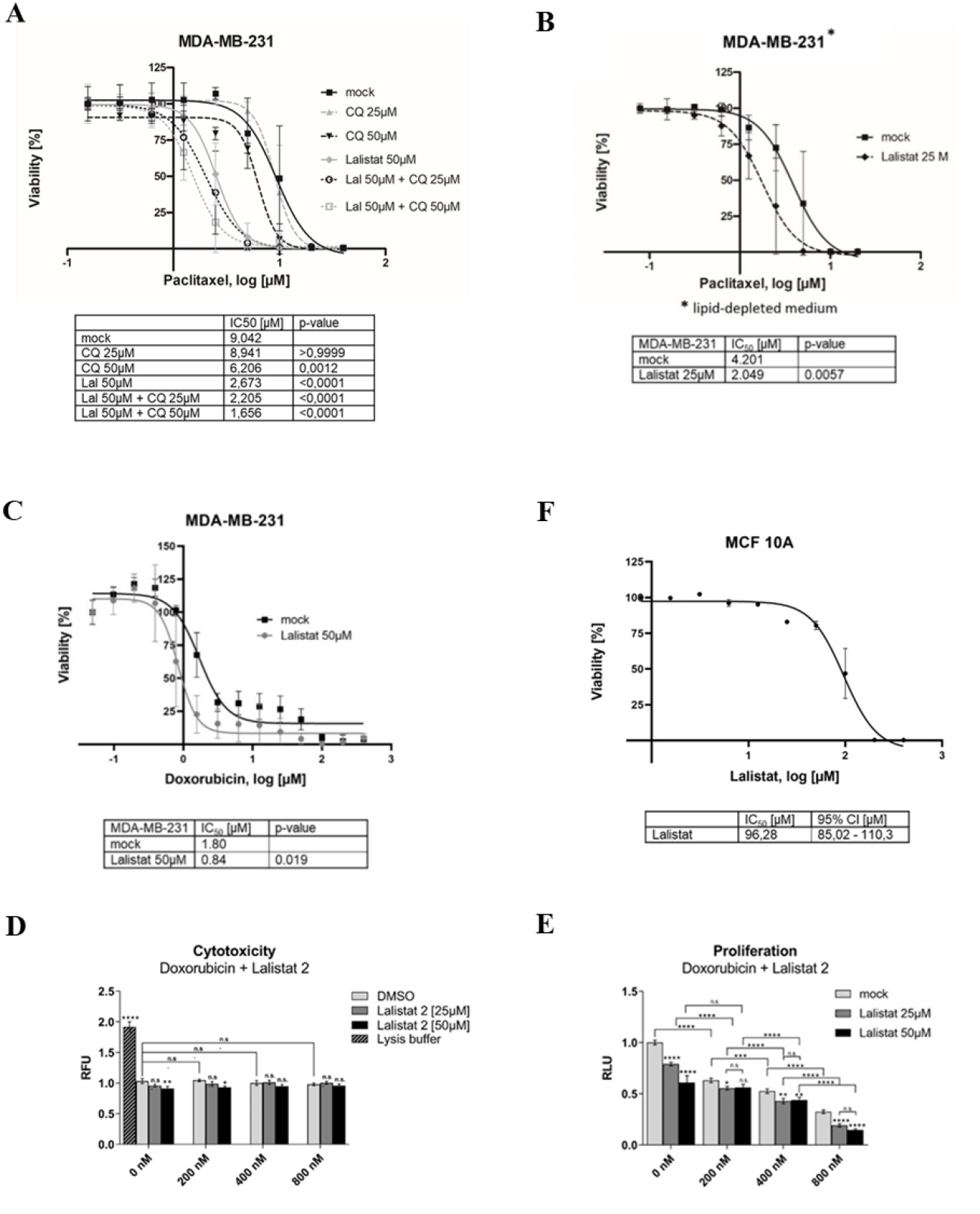
Lalistat shows high synergism with chemotherapeutics in TNBC. **(A, B)** MDA-MB-231 cells, incubated in lipoprotein-depleted medium in (B), were pretreated with chloroquine (A), Lalistat alone (A, B) or combined (A) in indicated concentrations for three days followed by combined incubation with paclitaxel in dilution series for further three days. Cell viability as a function of Paclitaxel concentration was evaluated by CellTiter-Glo^®^. Mock corresponds to solvents of Lalistat (A, B) and chloroquine in form of DMSO and H_2_O (A) in corresponding concentrations. The graph shows the mean +/-SD, n=12. In the table below IC50-values of Paclitaxel were compared. P-values ≤0.05 show significant difference compared to mock. All values were set in relation to viability of untreated cells. **(C, D, E)** MDA-MB-231 cells were incubated with Lalistat in 25 μM (D, E), 50 μM (C, D, E) or 0,05% DMSO (C, D, E) in combination with Doxorubicinin in dilutions series (C) or in different non-toxic concentrations (D, E). (C) Cell viability as a function of Doxorubicin concentration was evaluated by CellTiter-Glo^®^. In the table below IC_50_-values of Doxorubicin were compared. P-values ≤0.05 show significant difference compared to mock. All values were set in relation to viability of untreated cells. Mock corresponds to solvents of Lalistat. (D) Toxicity of Doxorubicin in lower concentrations in combination with DMSO or Lalistat was excluded by toxicity test. The cell toxicity was determined in form of relative fluorescence by using CellTox™ Green Cytotoxicity Assay. The positive control corresponds to lysis solution included in the assay. All values received were set in relation to blank-mean in form of untreated cells. (E) Cell proliferation was determined by CellTiter-Glo^®^ assay. All values received (D, E) were set in relation to blank-mean in form of untreated cells. The graphs show the mean +/-SD, n=8 (C) and n=4 (D, E). **(F)** IC_50_ -curve of Lalistat in MCF 10A: MCF 10A-cells were treated with Lalistat in dilution series for six days and evaluated for viability by CellTiter-Glo. The graph shows the mean +/-SD, n=4. The table shows IC_50_ of Lalistat and 95% confidence interval. Statistical analysis was performed by using one-way Anova. * = p ≤ 0,05; ** = p ≤ 0.01 *** = p ≤ 0.001; **** = p ≤ 0.0001

## Notes

### Competing Interest Statement

The authors have declared no competing interest.

